# A correlational study between microstructural, macrostructural and functional age-related changes in the human visual cortex

**DOI:** 10.1101/2022.03.17.484772

**Authors:** Sahar Rahimi Malakshan, Farveh Daneshvarfard, Hamid Abrishami Moghaddam

## Abstract

Age-related changes in the human brain can be investigated from either structural or functional perspectives. Analysis of structural and functional age-related changes throughout the lifespan may help to understand the normal brain development process and monitor the structural and functional pathology of the brain. This study, combining dedicated electroencephalography (EEG) and magnetic resonance imaging (MRI) approaches in adults (20-78 years), highlights the complex relationship between micro/macrostructural properties and the functional responses to visual stimuli. Here, we aimed to relate age-related changes of the latency of visual evoked potentials (VEPs) to micro/macrostructural indexes and find any correlation between micro/macrostructural features, as well. We studied age-related structural changes in the brain, by using the MRI and diffusion-weighted imaging (DWI) as preferred imaging methods for extracting brain macrostructural parameters such as the cortical thickness, surface area, folding and curvature index, gray matter volume, and microstructural parameters such as mean diffusivity (MD), radial diffusivity (RD) and axial diffusivity (AD). All the mentioned features were significantly correlated with age in V1 and V2 regions of the visual cortex. Furthermore, we highlighted, negative correlations between structural features extracted from T1-weighted images and DWI. The latency and amplitude of the three dominants peaks (C1, P1, N1) of the VEP were considered as the brain functional features to be examined for correlation with age and structural features of the corresponding age. We observed significant correlations between mean C1 latency and GM volume averaged in V1 and V2. In hierarchical models, the structural index did not contributed to significant additional variance in the C1 latency after accounting for the variance associated with age. However, the age explained significant additional variance in the model after accounting for the variance associated with the structural feature.

## 1. Introduction

The human brain undergoes extensive structural and functional changes from infancy to adulthood. Age-related changes of the human brain can be explored from two structural and functional perspectives. While the structural approach concerns the geometrical, dimensional and anatomical changes of the brain; alterations in behavior and neurological responses to different stimuli during the aging are considered in the functional approach. The functional investigation is mainly based on evaluating age-related changes in electrophysiological activity or metabolism of the brain. Several functionalities such as the speed of problem solving as well as the memory and learning degrade as a normal part of aging.

The cerebral cortex which constitutes more than half of the brain volume, plays a prominent role in phenomena like episodic memory, movement, thought, perception and attention. It consists of a wide variety of different cell types that have distinct genetic, functional and structural characteristics (1) (2). Investigating structural changes in both the cerebral cortex and white matter (WM) is of great interest. A steady increase in WM volume has been observed from birth and even beyond adolescence into middle age, while it continuously declines afterward. On the other hand, gray matter (GM) volume follows an inverted U-shaped trajectory peaking at around first years after birth and continuously declines later (3) (4). There are several mechanisms underlying the loss of GM volume and density such as the shrinkage, synaptic pruning (5) and cell loss (6), as well as the continued myelination which is extended to late adolescence (7). The prolonged procedure of myelination is concerned with cortical responsibilities. The complex phenomena such as the high level cognitive functions take more time to develop and require different levels of myelination (8).

Quantitative analysis of magnetic resonance images (MRIs) is important in identifying and evaluating anatomical brain changes, neurodegenerative and brain disorders. Some morphometric parameters of the brain cortex have been extracted and considered in the study of structural development such as the thickness, surface area, volume, curvature and folding of the cortex in different brain regions (9)(10)(11). Structural features such as the volume and weight of the brain diminish in elderly subjects (12) which are related to the shrinking of the cells (13). Storsve et al. (14) indicated an average annual percentage reduction of 0.35 for the cortical thickness of subjects aged between 23 and 87 years. Several other studies proved a similar declining procedure by aging (15) (16) (17) (14) (18). Aging-related reduction of the cortical thickness is also a consequence of cellular alternation like changing in the myelination or synaptic pruning (19) (20). Storsve et al. (14) further reported the annual 0.19 percent reduction in surface area of the cortex which is consistent with other studies (21) (22) (23) (24) (25). During aging, the surface area is reduced due to the reduction of dendritic size and complexity (26) (27). The cortical surface area and folding are essential parameters of cognitive development, optimizing the functional organization. Magnotta et al. (15) reported that by aging, the gyri become more steeply curved and sulci widen (28) which lead to declining of the local gyrification index (LGI) (29) (30) (31) (32). This reduction of LGI is parallel to the decline in surface area.

Brain changes due to aging can be further investigated using microscopic information provided by diffusion tensor imaging (DTI) (33) (34). The measures commonly used in diffusion MRI (dMRI) include mean diffusivity (MD), fractional anisotropy (FA), axial diffusivity (AD) and radial diffusivity (RD). These parameters are sensitive to the microstructures of the neurons such as the myelination, axon density, and axon coherence. MD represents the average magnitude of water displacement by diffusion, while FA measures the anisotropic diffusion and is sensitive to alignment of water-restricting barriers (35) (36). AD reflects the direction of maximum diffusion which quantifies diffusion parallel to the axonal fiber, and RD is supposed to increase with demyelination and reduction in axon density (36). There is evidence that a decline in the length and number of myelinated fibers as well as a breakdown in axon density will happen in normal aging (37) (38).

The supragranular layers of the cortex (layers I, II, and III from the six layers) contain predominantly neurons that project their axons to associative or commissural areas (39). This provides a strong background that myelin exists even in the GM of the cortex. In general, GM is mostly composed of dendrites and neuronal cell bodies as well as myelinated axons. In fact, there are little explorations about myelination in GM compared to WM. Uddin et al. (40) studied the diffusion parameters of subcortical GM or some cortical regions like anterior and posterior cingulate and insular cortex. It has been supposed that the increase of MD in GM reflects the breakdown of microstructural barriers to diffusion, indicating probable volumetric alterations (41). Therefore, GM’s MD could be a valuable measure of early GM cell damage. DTI measures were particularly aimed at highly anisotropic structures. In spite of the fact that most parts of the cortex contain GM and are relatively isotropic, some studies assessed DTI parameters in GM structures due to the presence of ferritin bound iron which could influence the DTI measurements (42) (43). Due to the fact that iron accumulates during the individual’s life, the study of DTI parameters in older ages could be beneficial (44). Glasser et al. (106) proved that the T1 and T2-weighted imaging are sensitive to microstructural features and changes in these images can be related to tissue properties such as the cell density and iron concentration that influence water mobility (92). Therefore, investigating the associations between the features from T1-weighted images and DTI would broaden our perspective. For instance, the results revealed in (45) showed that MD, AD and RD are correlated with WM volume.

Functional analysis of age-related changes can be performed by using event-related potentials (ERPs). Characteristics of the ERPs’ components (i.e. positive and negative deflections) represent the cortical activity and cognitive brain functions in response to a specific stimulus (46). While there is an increase in the peak amplitude and a decrease in peak latency of the responses at the first years of life, during aging the latency increases and the amplitude declines in some cases and is stable in others (47) (48) (49). In general, the first ERP components when elicited by visual stimuli are correlated to attention, processing of color, shape, rotation and degree of attention which are considered as simple visual analysis. On the other hand, the last components which are associated with discrimination tasks revealing more complex cognitive processes (50). The visual C1, generated in the striate visual cortex, is considered to reflect the processing of elementary visual features (the “analysis” stage) (51). Similarly, P1 and N1 peaks, are generated largely in the primary and secondary visual cortex (V1 and V2) as a response to the central region of the visual field that receives input from the lateral geniculate nucleus in the thalamus (52) (53) (54) (55). Nearly, all visual information reach the cortex via V1 as the largest and most important visual cortical area. V2 as the second major area in the visual cortex, and the first region within the visual association area, receives strong feedforward connections from V1 and sends strong projections to other secondary visual cortices. Myelination as a maturational process is the strong ground for these developmental changes. Increasing the latency and decreasing the conduction velocity during the aging is related to demyelination as a biochemical and neurophysiological change of the brain (56).

Although several studies investigated the structural and functional age-related changes of the human brain separately, there are only a few studies focusing on the joint structural and functional changes of the brain during aging. The focus on using computational methods to extract the multivariate biomarkers, which are informative indicators for brain aging, pathologies and the relation between different modalities, is increasing. Using informative features, multivariate biomarkers and merging information of all individual markers to assess the correlation between modalities are expected to have more reliable and accurate results. MRI-based studies have shown that brain aging is associated with distinct structural changes in the WM and GM. However, the relation between these changes and brain function is still poorly understood. In (57), the relationship between GM volume and EEG power was investigated showing that the reduction of GM (i.e. reflection of neurons or neuropil’s elimination) gives rise to the reduction in amplitude of EEG activity and absolute EEG power. In (58), the correlation between structure and function of the brain was investigated with increasing age, attempting to relate the gamma frequency band response of the primary visual cortex (V1) to age-related structural changes. While significant correlations were observed between the age and the gamma-band response as well as cortical features of pericalcarine and cuneus, hierarchical regression analysis failed to show unique effects of age or structural measure of cuneus on transient gamma peak frequency. Recently, Price et al. (59) related the latency of magnetic visual and auditory evoked fields (MEG) to age-related changes in the GM volume and WM microstructure (by DTI) across the whole brain. Their results confirmed that while the functional properties of the visual responses were related to the microstructural properties of the WM, properties of auditory responses were related to the cortical GM and efficiency of local cortical processing.

In the current study, MR images and EEG signals were used for modeling structural and functional changes of the brain, respectively. Furthermore, DTI was used for extracting microstructural parameters to be investigated for correlations with the functional and structural features of T1-weighted MR images. A challenging question of “how structural changes in the brain lead to enhancement or degradation of the brain functions during the human life?” has encouraged us to study joint structural and functional changes of the visual cortex with increasing age. It has been shown previously, that WM microstructure in the optic radiation (connecting thalamus to the visual cortex) partially mediates age-related constant delay of the visually evoked fields which leads to increased transmission time for the arrival of information communicated from LGN to V1 (59). Apart from these associations that are related to WM demyelination occurred by aging, we hypothesized that age-related delays may also depend on some factors affecting the speed of processing after the visual information reaches the cortex. Specifically, the cortical GM microstructure would reflect the age-related neural conduction slowing in the VEPs. Therefore, we extracted the amplitude and latency of the components at occipital electrodes as the functional features, as well as several structural features of the visual cortex from MRI and DTI in a broad range of age (20-78 years). It is worth mentioning that similar relationships have been investigated between microstructure measures of cortical gray matter and auditory evoked responses in adults (59), though these effects disappeared after adjusting for the effect of age. DTI parameters seem to have relative sensitivity to neural changes and provide us with a wealth of information about brain aging. DTI might be a practical tool for following the visual cortex microstructural changes and reveal the existence of association with either functional or other macrostructural changes. To the best of our knowledge, this is the first study on investigating joint functional and structural age-related changes by extracting features from these three different modalities. Here, we aimed at presenting the functional and structural age-related changes of the visual cortex, as well as investigating the correlations between the structure and function at this specific brain region.

## 2. Materials and methods

The specific approach, proposed in the current study, for joint structural-functional analysis of the brain aging in the adults’ visual brain cortex can be presented as Fig 1. In the following sections, these procedures and results will be described in detail.

**Figure 1.**
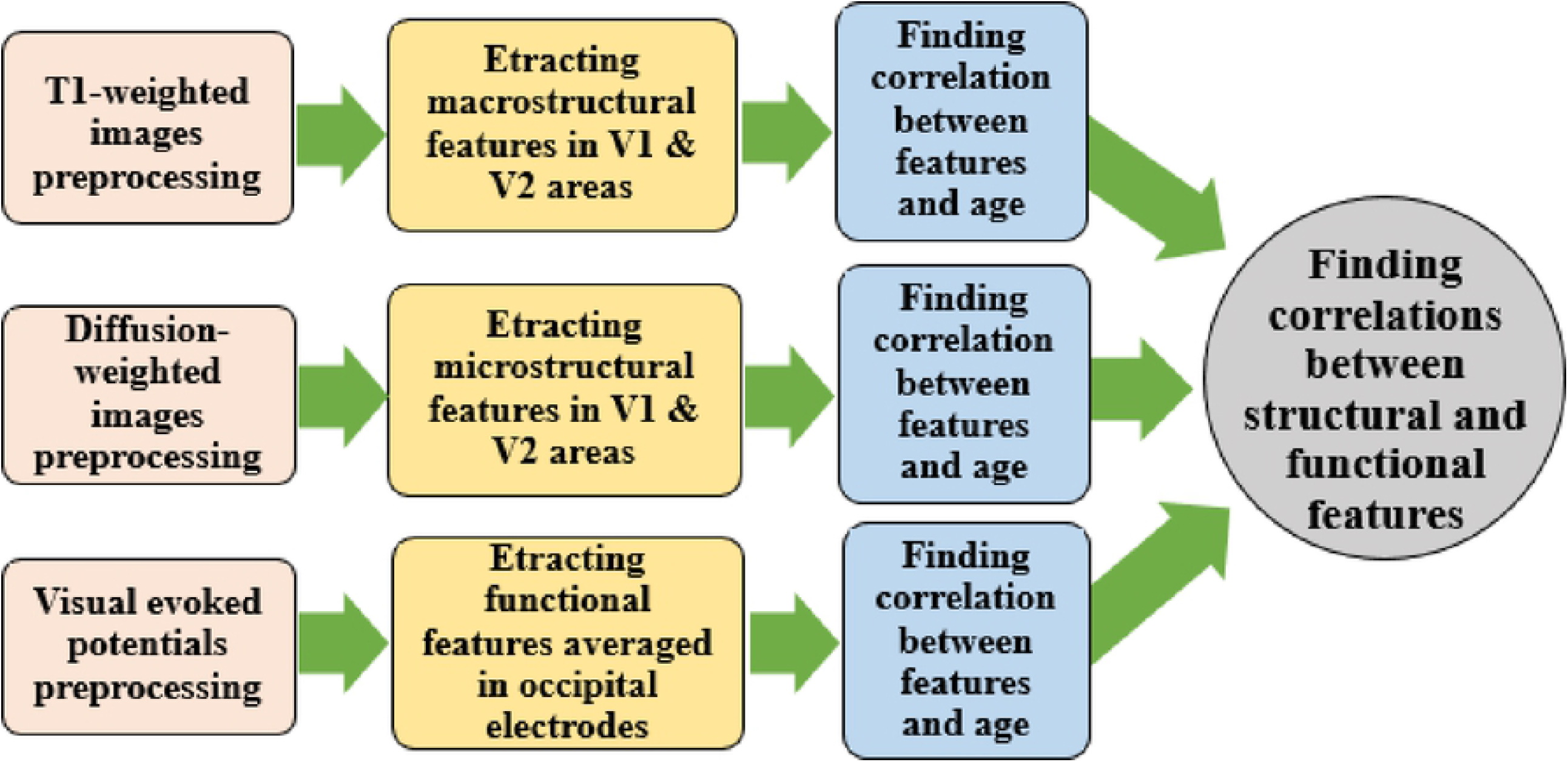
The specific approach proposed for joint structural-functional analysis of the visual cortex alterations due to aging.

### 2.1 Functional data

#### 2.1.1 Participants

The functional dataset includes the EEG signals recorded from 45 healthy adults with an age range of 20 to 78 years (recruited in (51)) in response to the visual and auditory stimuli. The neurobiological, psychiatric and medical history of the participants was checked. All the participants had normal or near to normal vision. Details related to the dataset and subjects’ information are presented in the Auditory-Visual Attention Shift Study of HeadIT Data Repository (HDR)^1^. Four subjects were exempted from our study due to the lack of information regarding their age.

#### 2.1.2 Stimuli

The procedure of the original recording consisted of both visual and auditory stimulations. The visual (target and non-target) and auditory (target and non-target) stimuli were presented in focus attention and shift attention blocks. In the shift blocks, not applied in the current study, subjects shifted their attention between the visual and auditory modality based on the presentation of ‘Look’ and ‘Hear’ cues. In the focus blocks, they attended the cue, which was instructed prior to the start of the block. For each subject, one focus auditory block (FAB) and one focus visual block (FVB) were presented, followed by 12 shift blocks and another 6 blocks including random FAB and FVB. The order of the focus blocks was randomized across subjects. In the current study, we used non-attended non-target visual stimuli obtained in focus auditory conditions to reduce the effect of attention. Since the facilitation and inhibition are related strongly to attention or its withdrawal, avoiding the attentional confounds helps in a better understanding of age-related changes regarding the sensory domain. The visual target and non-target stimuli were light and dark blue squares (8.4 cm^2^), respectively. They subtended 3.3 degrees of visual angle and were presented for 100 ms on a light gray background of a computer (51). Subjects were asked to press a button upon detection of attended target stimuli during the presentation of different blocks.

#### 2.1.3 EEG recording and preprocessing

The EEG signals were recorded from 33 scalp electrodes (FP1, FPz, FP2, AF3, AF4, F7, F8, F3, Fz, F4, FC1, FC2, FC5, FC6, T7, T8, C3, Cz, C4, CP1, CP2, CP5, CP6, P3, Pz, P4, P7, P8, PO3, PO4, O1, Oz, O2) based on the international 10-20 system. The signals were amplified by 10 and filtered at 0.1-60 Hz. Reference of the recordings was right mastoid and the recordings were re-referenced to the average of the left and right mastoid. Independent component analysis (ICA) has been used to eliminating various kinds of artifacts. The signals were segmented into epochs from 200 ms pre-to 600 ms post-stimulus onset for the non-target visual stimulation in FABs. Baseline correction was performed according to 200 ms before the beginning of the stimulation and body movement artifacts, as well as false-alarm button press, were excluded from the epochs. The epochs were then averaged for each subject to obtain ERPs at each electrode. In our study, all of the preprocessing steps have been done by EEGLAB.

#### 2.1.4 ERPs components

By taking simple visual stimuli and grand average source localization into account, C1, P1 and N1 peak were extracted for each subject at three occipital electrodes (O1, O2 and Oz). These values were then averaged across the channels to obtain one functional feature for each component’s amplitude and one for each component’s latency. The ERP peak search windows were defined by visual inspection of the grand average waveforms according to the paper that employs the same dataset as our study (51). The features were then examined for the age and structure-related dependencies.

### 2.2 Structural dataset

#### 2.2.1 MRI and DWI datasets

To develop a conjoint model of functional and structural brain changes, a structural dataset corresponding to the same age-range of the functional data is required. Ideally, the structural data acquired from the same subjects recruited for functional data acquisition are ideal for the joint analysis. However, due to the unavailability of structural data from the same subjects, we selected T1-weighted MR images from the Imperial College London repository (IXI dataset^2^) which contains almost 600 normal healthy subjects with the age range of 20-86 years. MRI data were collected from three different scanners (at three different hospitals in London), two of which were 1.5 T and one was 3 T scanner. Details of the IXI data and scan parameters are available at: (http://brain-development.org/ixi-dataset/).

Furthermore, the DWI dataset released by the Cambridge Centre for Ageing and Neuroscience (CAMCAN)^3^ was employed, which includes approximately 600 normal subjects aged between 18-87 years (62) (63). DWIs were acquired with a twice-refocused spin-echo sequence, with 30 diffusion gradient directions for each of the two b-values: 1000 and 2000 s/mm2, plus three images acquired with a b-value of zero. The other parameters were as follows: Echo Time = 104 ms, repetition time = 9100 ms, voxel size = 2 mm isotropic, FOV = 192 mm × 192 mm, 66 axial slices, number of averages = 1. The images were processed for extracting DTI parameters (FA, MD, AD and RD). In our study, we used the data of one person at each age (i.e. each year) to extract the features. This leads to considering 59 subjects (20-78 years) from each of the MRI and DTI datasets.

#### 2.2.2 Structural analysis

We processed MR images using the default recon-all preprocessing pipeline of FreeSurfer^4^ (Center for Biomedical Imaging, Charlestown, MA). This procedure consists of brain extraction, motion correction, intensity normalization, registration, segmentation, tessellation, generation of white and pial surfaces, surface topology correction, spherical mapping, registration to the FsAverage surface based on a measure of surface shape, and finally cortical parcellation mapping. In the next step, we extracted various features including the surface area, GM volume, folding index, curvature index and average cortical thickness from MR images in V1 and V2 cortical regions which are responsible for processing the presented simple visual stimuli.

Cortical thickness is commonly defined as the distance between the white surface (the border between WM and GM) and the pial surface (the border between GM and CSF) (64) (16). The cortical surface area is calculated by summing the areas of all triangles that form the WM/GM boundary within a specific region (65). While the surface area and thickness of the whole cortex or a region of interest (ROI) are known, the cortical volume can be obtained by a simple multiplication (66). Considering that *k*_*1*_ and *k*_*2*_ are the maximum and minimum curvature respectively at any point on a surface cortex, the remaining features can be calculated in each ROI by a combination these two factors (equations 1, 2) (67) (68).

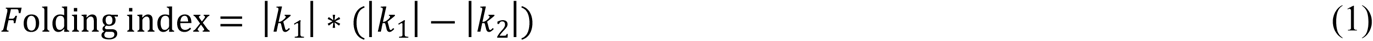

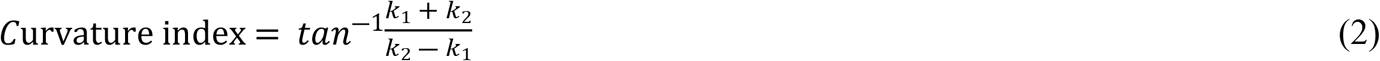

For DWIs, eddy current distortions and subject motion artifacts were corrected using an affine alignment of each diffusion-weighted image to the b = 0 image using FMRIB diffusion toolbox^5^ (FSL 4.0). DTI parameters including AD, RD, MD and FA were extracted for V1 and V2 regions using modified code from ExploreDTI (A. Leemans, Utrecht, Netherlands) (69). All the structural features were averaged across V1 and V2 to be investigated for the age and function-related dependencies. Furthermore, linear regressions were conducted to investigate the correlations between the structural features of T1-weighted images and DTIs.

### 2.3 Joint functional-structural analysis

To investigate correlations between the function and structure, representative functional and structural features are required for each specific age. Therefore, the features were extracted for one subject at each year in the functional as well as MRI and DWI datasets. Linear regressions were performed to investigate the correlation between the functional and structural features. Furthermore, hierarchical regressions were conducted to examine the unique influence of age and structural changes on the functional changes. All the statistical analyses were carried out using IBM SPSS Statistics.

## 3. Results

### 3.1 Functional analysis of age-related changes

Grand average ERPs, for young (20-40 years) and old (65-78 years) groups, averaged across occipital electrodes are presented in Fig. 2 to show the difference between these two groups. Despite considering the changes in latency and amplitude, Fig. 2 displays the appearance of P1 in the form of a double peak instead of a single one, occured by aging. As Fig. 3 shows, there is a significant positive correlation between the age and the latency of C1 (p = 0.009, r = 0.384) averaged in three occipital electrodes. However, no significant changes were observed for the amplitudes of C1 with increasing of age. Partial correlations were conducted to examine the strength of age-related correlation after controlling for the effect of gender and the result was still significant. Furthermore, there were no significant correlations between the latency and amplitude of other components (P1 and N1) with age.

**Figure 2.**
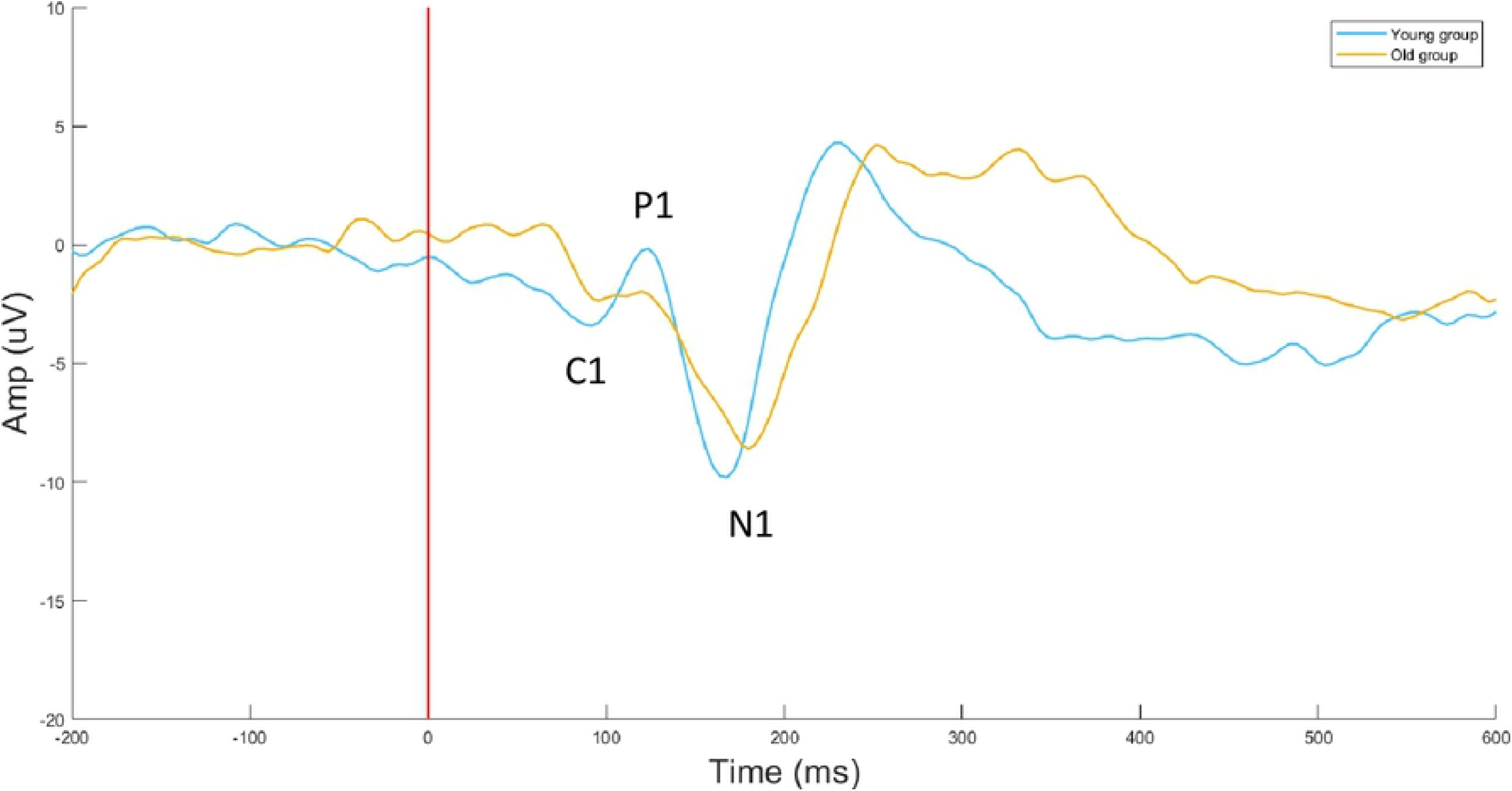
Grand-average ERPs for young (20-40 years old) and old (65-78 years old) groups, averaged across occipital electrodes (O1, Oz, O2).

**Figure 3.**
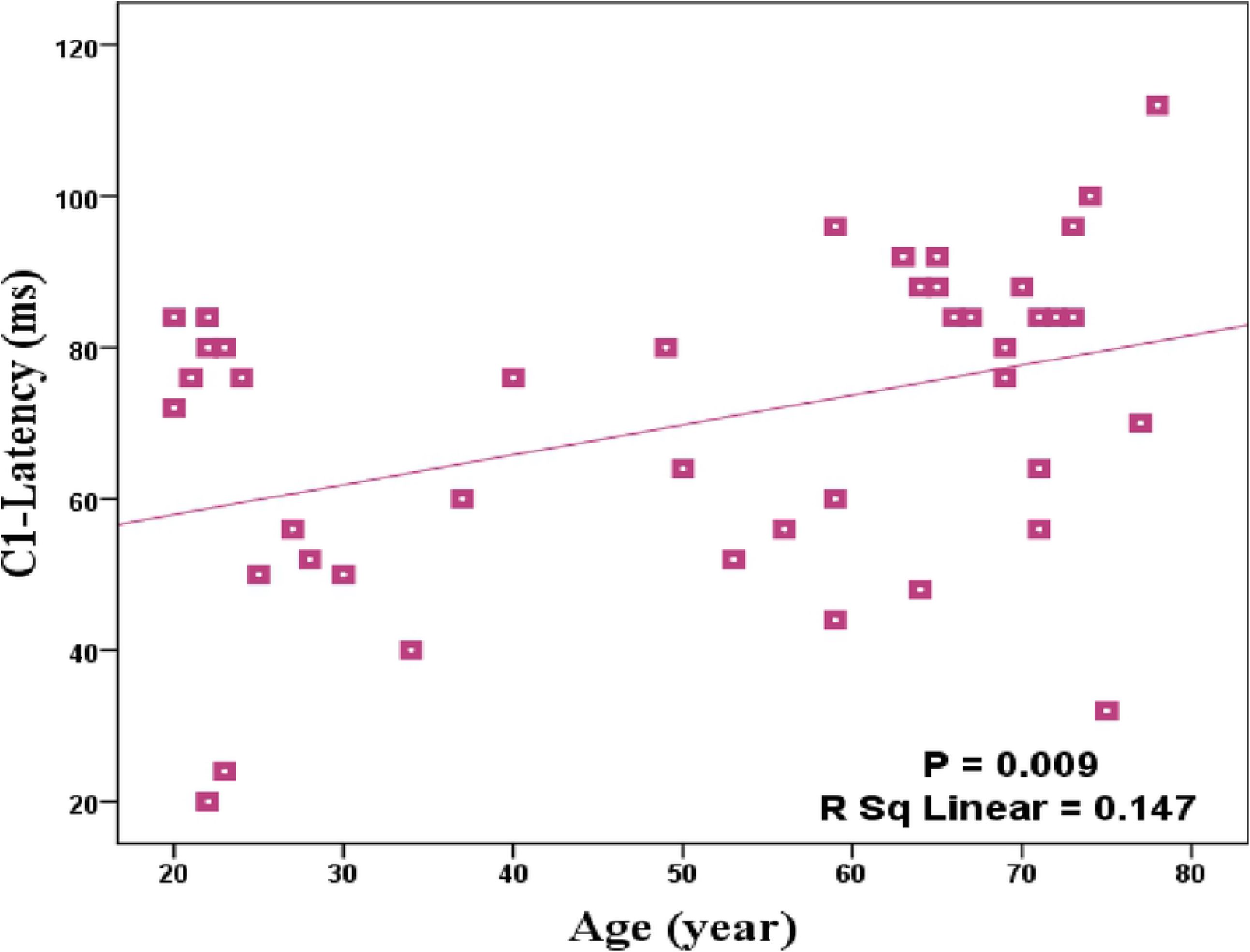
Significant age-related changes in C1 latency averaged in O1, O2 and Oz electrodes

### 3.2 Structural analysis of age-related changes

Significant negative correlations were observed between the age and surface area (p < 0.001, r = −0.55), cortical thickness (P < 0.001, r = −0.611), volume (p < 0.001, r = −0.587), folding (p < 0.001, r = −0.570) and curvature index (p = 0.003, r = −0.367) averaged in V1 and V2 regions (Fig. 4). All the results were significant even after partial correlation analysis which investigated the effect of age on the structural index while correcting for the effect of gender. While analyzing the DWIs, significant positive correlations were observed between the age and MD (P < 0.001, r = 0.445), RD (p = 0.002, r = 0.382) and AD (p = 0.001, r = 0.418) averaged in V1 and V2 regions, but not for the FA (Fig. 5). The results were still significant after controlling the gender effect. Moreover, we employed quadratic models to explore the age-related changes across the human lifespan that is shown for MD, RD and AD features in Fig. 5. The significant results indicate the higher *R*^2^ value for quadratic in comparison to the linear model.

**Figure 4.**
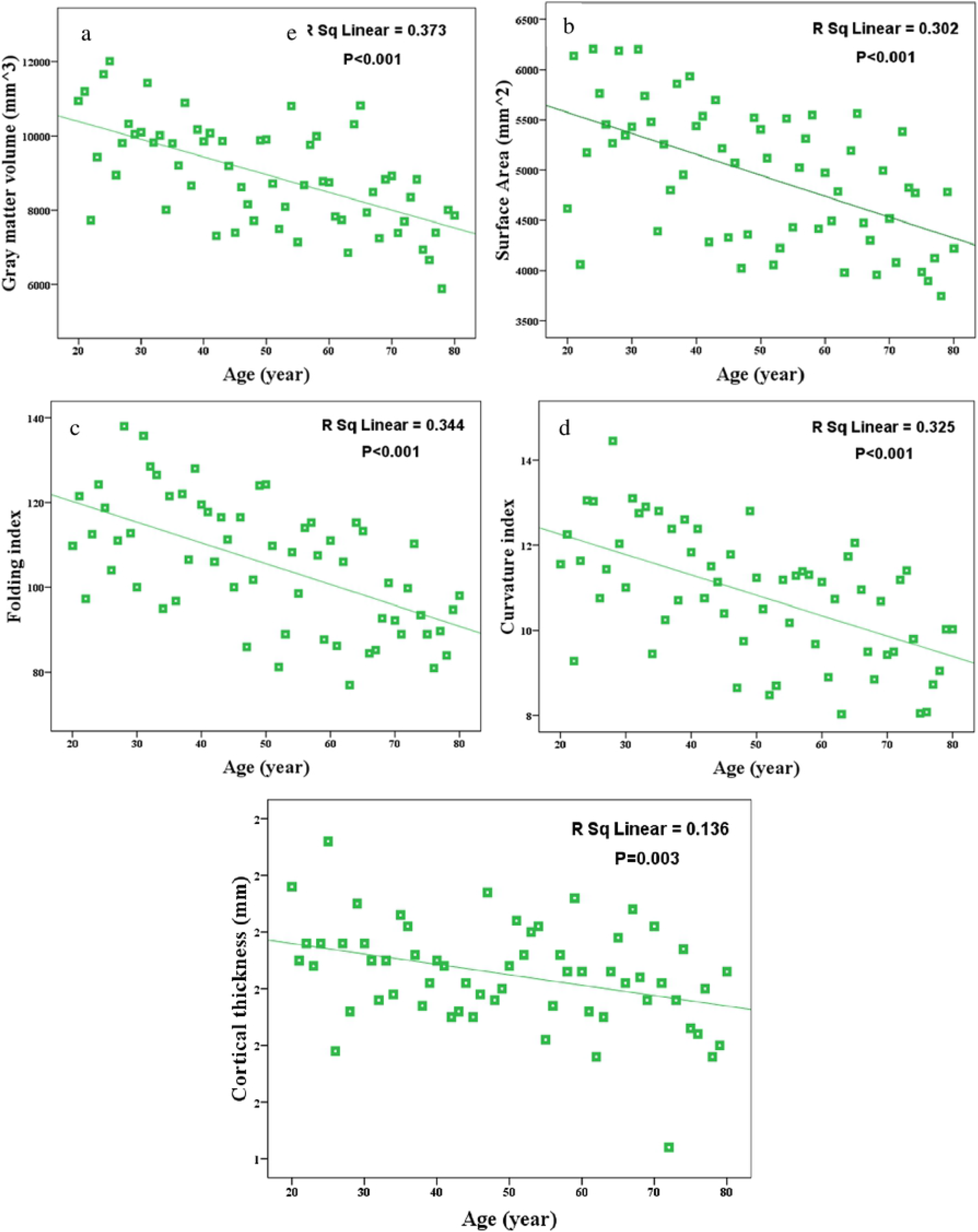
Significant age-related changes in a) surface area, b) GM volume, c) folding index, d) curvature index and e) mean cortical thickness averaged in V1 and V2.

**Figure 5.**
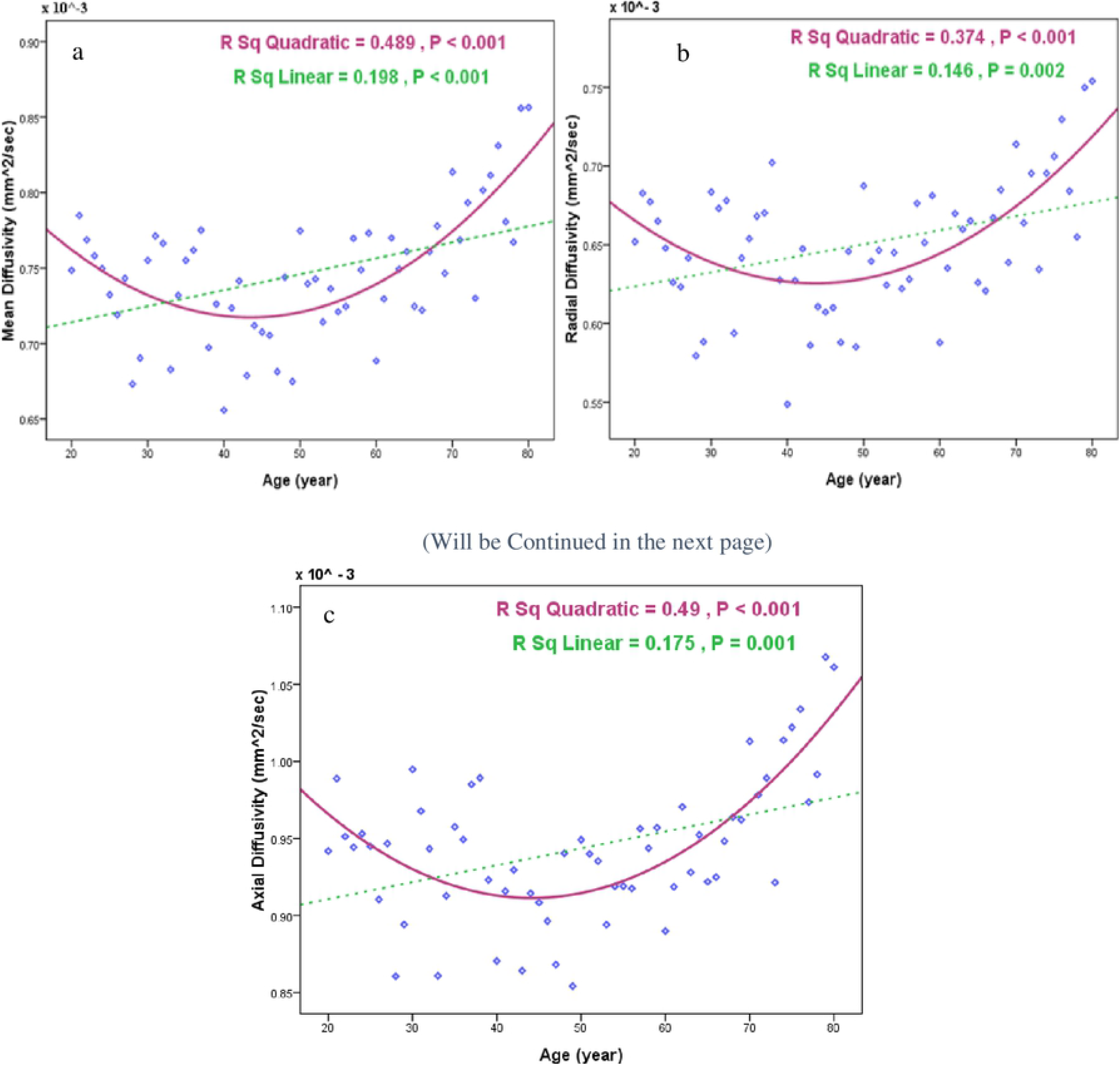
Significant quadratic age-related changes fit better in comparison to the linear model in a) MD, b) RD and c) AD averaged in V1 and V2.

### 3.3 Structure-function correlations

Significant negative correlations were observed between the C1 latency averaged in three occipital electrodes and GM volume (p = 0.048, r = −0.377) averaged in V1 and V2 regions (Fig. 6). We performed hierarchical regressions to investigate the unique effects of age and GM volume on the C1 latency. The full regression model (age and GM volume) predicted 51.6% of the variance in C1 latency (p < 0.001). Added first, age predicted 51.4% of the variance (p < 0.001). Including the GM volume explained a non-significant 0.2% of the additional variance in the model (p = 0.753). Added first, GM volume predicted 14.2% of the variance (p = 0.048). Including age explained a significant 37.4% of additional variance in the model (p < 0.001).

**Figure 6.**
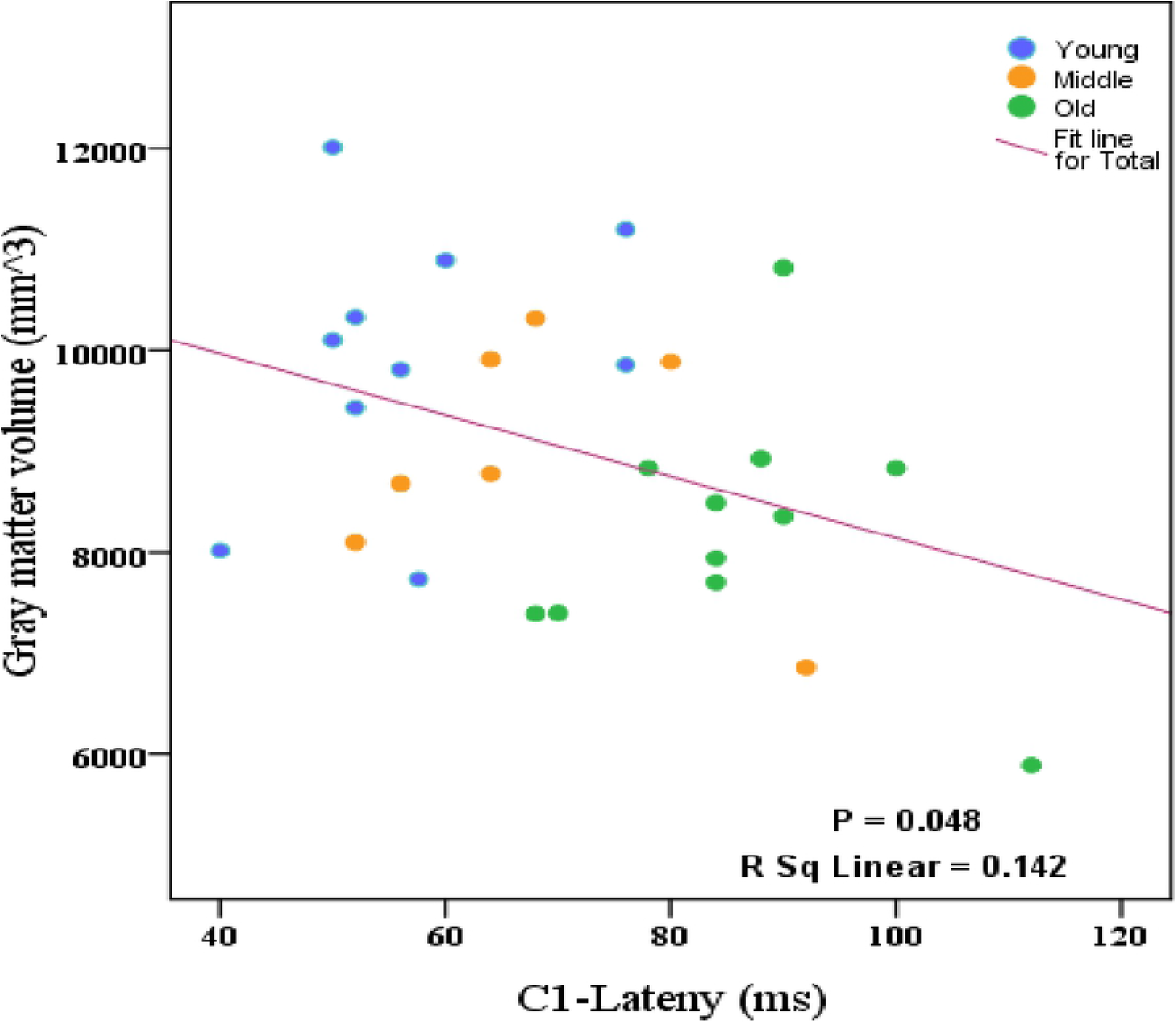
Significant correlations between mean C1 latency and GM volume averaged in V1 and V2; for young (20-40 years), middle (40-65 years) and old group (65-78 years).

In general, by applying hierarchical regression analyses to examine the relative contributions of age and cortical feature, GM volume, to the latency of C1 component in VEPs; we see that if structural feature was added first, both structural feature and age almost predict the significant variance in latency of the visual responses. However, after removing variance in the visual response latency associated with age, cortical characteristics accounted for no additional variance. As observed from the results, the correlation between the structural and functional features was not significant after removing the effect of age.

### 3.4 Structure-structure correlations

Table 1 presents the results of the negative correlations between the structural measures extracted from T1-weighted images and microstructural features computed from DTIs. Although significant negative correlations were observed between some features, the results were not significant after removing the effect of age.

**Table 1.** P-values and correlation coefficients for negative correlations between structural features extracted from T1-weighted images and DTI.

| Structural features extracted from T1-weighted images<br>Structural features extracted from DTI |  | Surface area | Gray matter volume | Folding index | Curvature index | Average cortical thickness |
| --- | --- | --- | --- | --- | --- | --- |
| Mean diffusivity (MD) | r | -0.287 | -0.244 | -0.348 | -0.375 | -0.132 |
|  | p | **0.025** | 0.059 | **0.006** | **0.003** | 0.312 |
| Radial diffusivity (RD) | r | -0.281 | -0.240 | -0.342 | -0.382 | -0.142 |
|  | p | **0.028** | 0.062 | **0.007** | **0.002** | 0.274 |
| Axial diffusivity (AD) | r | -0.253 | -0.228 | -0.341 | -0.342 | -0.149 |
|  | p | **0.049** | 0.077 | **0.007** | **0.007** | 0.251 |

## 4. Discussion

Our findings showed several associations between structural changes, electrophysiological function, and age, which present a biophysical model of age-related changes in the visual area of the human brain. The results presented regionally specific changes due to aging and may thus be informative for a better understanding of physiological processes of age-related decline. Here, structural features of the visual areas were extracted from T1-weighted images and functional features were extracted from the visual cortical responses of EEG signals. Moreover, in this study, DTI microstructural features of the specific regions including V1 and V2 in the visual cortex were extracted and correlated with the age, functional and structural features. Although significant correlations were observed between the structure and function, and also between the structural features of the T1-weighted images and DTIs, the correlations were not significant after controlling for the effect of age. To be more specific, although the GM volume was significantly (negatively) correlated with C1 latency, hierarchical regressions revealed that these correlations were not significant after removing the effect of age. Therefore, the increase in the C1 latency with increasing age is not likely the result of alteration in the structural features of the visual area. Rather the C1 latency, structural features and age may be related indices of the visual cortex aging. It is worth mentioning that age explained significant additional variance in the model after accounting for the variance associated with the structural feature, which in turn give rise to the idea that age, apart from the effect of the structural feature, contains biological underpinnings of age-related changes that occurred in V1 and V2 areas such as demyelination, fiber loss and degeneration.

Similar to our result, Stothart et al. (79) reported the appearance of P1 in the form of a double peak instead of a single one by aging which can be due to de-synchronization of cortical sources. De-synchronization was confirmed by results showing that activation in older subjects changed in comparison to younger adults. In the old group, activation distributes to frontal areas indicating a Posterior Anterior Shift in Aging (PASA) functional activity (70). Most of the VEPs findings reported age-related diminished activity in visual cortices which is consistent with the findings of fMRI studies (71) (80). In this respect, they showed greater activation in frontal and parietal regions coincided with reduced activation in the occipital region (81) as a result of the compensation mechanism as well as a behavioral weakening in visual perceptual abilities occurred with aging. Decreases in occipital cortex activity by aging have been attributed to disorganized sensory processing in the ventral pathways. In fact, increasing age brings about dedifferentiation (a cellular process in which a differentiated cell loses its special form or function) of neural response in the ventral visual cortex (72).

Age-related changes we observed for MRI features are in accordance with previous studies, regarding the surface area, cortical thickness, cortical volume and folding index, both regionally (71) (58) (82) and globally (23) (73) (14) (74). For the folding index and curvature index, it is the first time that age-related changes are reported for V1 and V2 regions and their reduction pattern was in line with constant decrease of the similar measure (gyrification) during the adult’s lifespan (75). The reduction observed in the cortical thickness might be associated with synaptic pruning and related decrease of glial cells due to reduced metabolic requirements (76). The decline in GM volume by aging, which has a higher R-squared value in comparison to other features, is due to a combination of synaptic exuberance, pruning of the myriad connections and neuronal loss as a regressive and progressive event. Furthermore, during aging, the surface area is reduced due to the reduction of dendritic size and complexity (26) (77) (27). The process of atrophy in the cortex brings about widening and flattening of the sulcal regions resulting in a decreasing pattern in curvature index (80). Similarly, the cortical folding reduction would be a marker of brain atrophy in later decades of life which happens by increasing of age (78). The reason for age-related changes in the folding pattern (74) is not adequately clarified yet. It is only hypothesized that perinatal pruning with programmed cell death and reduction of the cell numbers and connectivity results in cortical gyrification reduction.

Regarding the DTI parameters, age-related changes in FA were not significant. The reason behind this observation is that this metric mainly reflects the degree of non-isotropic diffusion, while the cortex is mostly an isotropic environment (38). For MD, the age-related increase was shown regionally in pericalcarine (79). Our report of U-shaped paths across the age, for MD, is similar to Grydeland et al. (79) findings in some cortical areas such as the insula and isthmus of cingulate gyrus. U-shaped trajectories, indicate an accelerated myelination process until 30 years of age, followed by a period of relative stability, before a decrease in myelin content from the late 50s. The results accord with the seminal histology study by Yakovlev and Lecours (80), who reported an increase of myelinated fibers in the association cortices until the third decade and possibly beyond. Furthermore, there is only one study considering deep GM structures (43) which showed similar higher DTI values in the older rather than the younger group. MD suggested as an inverse measure of membrane density, signified cellular changes in GM which consists of cell bodies and is an independent measure of direction.

According to the study conducted by Glasser et al. (106), T1 and T2 weighted imaging are sensitive to microstructural features of the brain and changes in these images can be related to some tissue properties such as the cell density and iron concentration that influence water mobility (81). To date, there is no finding regarding the correlation between GM features in T1 weighted and DTI indices, particularly in the visual cortex. However, Pareek et al., (82) and Kochunov et al., (83) examined global features and showed that WM diffusion measures have a strong correlation with GM volume along the aging process. Similarly, in our study, the correlations between the DTI indices and some GM features were significant and elucidated how GM’s microstructural parameters have an association with GM’s macrostructural characteristics, during normal aging. However, they were not significant after controlling the effect of age.

Although this study and related works in the literature showed associations between certain brain structures and functions during aging, it has not been determined yet which is the cause and which is the consequence for these relationships. To sum up, despite strong evidence that visual object processing slows down with age, the exact origins of the aging effects, as well as the origin of the inter-individual differences within age groups (84), are still unclear. Apart from physiological changes occurring in the aging brain, it is possible that part of the age-related delays is due to factors affecting the speed of processing before visual information arrives the cortex. Particular reduced retinal illumination and degeneration of the eyes and visual function appears in the aging eye or the pathways till the information transforms to the cortex (85) (86). For example, age-related differences in visual acuity and contrast sensitivity could result in delayed processing speeds. Hence, maybe this would be the reason why the latency of C1 does not correlate with cortical related features independent of age.

However, we can find in Adibpour et al. (61) a relation between the conduction speed of the auditory P2 and GM maturation (longitudinal diffusivity) in the inferior frontal region but no relationship with DTI parameters alterations in acoustic radiations in infants. The strong ground related to these findings would be various developmental mechanisms that occurred in GM maturation at this age such as the myelination of intra-cortical fibers and the increase in synaptic density (89). Additionally, in visual stimulus, they only found an association between P1 speed and optic radiations independent of age. Therefore, a recent study indicates that structure-function relationships differ between the visual and auditory modalities (59). In this respect, the aging demyelination in cortical regions which can influence DTI indices is not as obvious and effective as myelination and its influence on DTI measures in infants’ age; hence, we can suggest that the visual system would be different in a structure-function relationship compared to the auditory modality. Another reason would be the importance of paying attention to cortical properties. For instance, subcortical diffusion measures are highly affected by localized changes in iron accumulation that are specific and related to the subcortical GM regions of the brain among older individuals (90) (91). Because, there is no measurement in the present study related to the visual cortex, the possibility that cortical differentiation of diffusion measures are associated with iron accumulation would be a hypothesis. So, we can propose that cortical myelination in the visual cortex is not an accurate marker and is not associated with the function of visual cortex, independent of age.

The existence of several microstructural components including the cellular membranes, axonal sheaths, and organelles in the human brain tissues, specifically in the cerebral cortex which is a complex microenvironment, leads to non-Gaussian diffusion of water molecules which is not limited to an anisotropic environment (92). In this situation, diffusion would be better characterized by a high order tensor diffusion model (93) called diffusion kurtosis imaging (DKI). In this regard, metrics such as the mean kurtosis (MK), radial kurtosis (RK) and axial kurtosis (AK) (94) (95) have been used in various studies and are sensitive to microstructure changes either in aging (96) (97) or disease (98) (99). Moreover, neurite orientation dispersion and density imaging (NODDI) have been proposed to indicate decent interpretation in areas of complex axonal or dendritic architecture. NODDI provides a sophisticated model of dMRI and supposes three distinct tissue compartments for water diffusion (intraneurite water, extraneurite water and cerebrospinal fluid) (100). NODDI changes have been reported in some studies related to cortical GM variations during aging (101), schizophrenia (102) and epilepsy (103). NODDI model fully explains the effects of free-water contamination, which is a considerably significant issue in aging (104). However, there is a lack of investigations and details regarding distributions of NODDI measures in the cortical GM which requires further studies.

In general, the MRI-dMRI studies might provide the foundation of the multifactorial changes in the structural GM volume and WM diffusion characteristics. In addition to the DTI parameters which present the structural markers of myelination, another MR-extracted feature is proposed which may reflect age-related changes resulted from the myelination. The ratio of T1/T2 weighted MRI signal intensity is supposed to improve the sensitivity to measure myelin-related signal intensity changes. This method provides the possibility to measure the myelination for both GM and WM regions; since it does not depend on the axon fiber anatomy compared to DTI (105). This measure has been used previously as an estimate of the myelin content in adults (105) (106) (79) and may be measured in future works to be correlated with the components’ latencies which are directly related to the myelination.

We should notice that the casual mechanism of joint structural and functional associations is still an interesting and not well-defined subject. The most crucial point regarding joint analysis is acquiring structural and functional data of the same subjects at the same time that is recommended to characterize the aging more accurately. Additionally, in joint structural-functional investigations, selecting structural and functional markers to be age predictive and have a correlation with each other is a challenging topic.

## Footnotes

1 https://headit.ucsd.edu/studies/8d6dbb78-a236-11e2-b5e7-0050563f2612

2 https://brain-development.org/ixi-dataset/

3 http://www.mrc-cbu.cam.ac.uk/datasets/camcan/

4 http://surfer.nmr.mgh.harvard.edu/

5 http://www.fmrib.ox.ac.uk/fsl/

## Notes

### Competing Interest Statement

The authors have declared no competing interest.

